# Boosting Protein-Protein Interaction Detection with AlphaFold Multimer and Transformers

**DOI:** 10.1101/2025.03.05.641631

**Authors:** Lena Connesson, Gabriel Krouk, Baldwin Dumortier

## Abstract

In 2021, DeepMind released AlphaFold 2, an AI-driven algorithm that revolutionized protein folding predictions by achieving error rates comparable to traditional experimental methods. Its remarkable performance has inspired researchers to extend its applications beyond its original purpose, leveraging AlphaFold 2 and its output features to tackle various unresolved biological challenges. In this paper, we discuss previous works that explored the repurposing of AlphaFold 2 for the detection of peptide-protein interactions (PPI). We propose a novel methodology where the outputs of a lightweight version of AlphaFold 2 Multimer are enhanced by a fine-tuned transformer encoder and benchmarked against classic machine learning algorithms. We study the performance of our approach on a curated version of Propedia, a peptide-protein dataset, and show that the boosting models significantly improve the detection of PPIs. Additionally, we provide HAPI (Hacky API), a freely available Python library designed to facilitate the seamless integration and use of AlphaFold 2 in computational workflows, further empowering the scientific community.

## 1 Introduction

Proteins are fundamental to cellular functions and play a crucial role as regulatory targets in medical and agronomy applications. These large biomolecules consist of chains of varying lengths, composed of combinations of the 20 naturally occurring amino acids, or residues. The three-dimensional structures of proteins, which are influenced by their amino acid sequences and post-translational modifications, determine their chemical and mechanical properties (Pabo [1983]). Traditionally, understanding the connection between a protein’s sequence and its structure was a major challenge in computational biology, often approached through physical modeling techniques (Leaver-Fay et al. [2011]). However, the advent of deep learning and the growth of 3D structure protein databases have significantly advanced this field, leading to groundbreaking developments (Min et al. [2017]).

A major advancement came with AlphaFold 2 (AF2)(Jumper et al. [2021]) and RoseTTAFold (Baek et al. [2021]), which addressed the protein folding problem using a multi-path transformer architecture. These models combine evolutionary data through Multiple Sequence Alignments (MSA) with graph embeddings of residue interactions, constrained by geometric properties, to improve protein structure predictions. AF2’s exceptional performance at the 14th Critical Assessment of Protein Structure Prediction (CASP14) demonstrated a significant leap forward, leading to a reassessment of the monomer folding challenge and underscoring the model’s impact. These models represent a significant leap forward, providing high-precision predictions of protein structures that have set a new standard in the field. Their success has catalyzed further research, inspiring the application of AI to a broader range of protein-related tasks. The remarkable performance of these models has not only revolutionized structural biology but has also opened up new avenues for leveraging AI in areas such as protein design, interaction prediction, and drug discovery.

Notably, one of the earliest and most impactful extensions of these models has been their adaptation to predict multimeric protein complexes. Researchers initially used creative workarounds to extend AF2’s capabilities to multimeric systems (Mirdita et al. [2022], Ko and Lee [2021], Tsaban et al. [2022], Bryant et al. [2022a]). This included modifying the original models and using innovative strategies to predict how different proteins interact within a complex.

Subsequent works, which focused on binder generation and enabled through hallucination or evolutionary strategies (Jeliazkov et al. [2023], Verkuil et al. [2022], Jendrusch et al. [2021], Goverde et al. [2023]), provided new extensions of AlphaFold2 able to score and predict protein-protein interaction. In particular, AlphaFold Initial Guess (AFIG, Bryant et al. [2022a]), or similar works by Bryant and Elofsson [2022], Bryant et al. [2022b], studied and improved speed of AF2 to score PPI by scooping the MSA clustering tools and by providing the expected structure template to the neural network part of AlphaFold2.

Later, recognizing the limitations of these workarounds, DeepMind released AlphaFold Multimer (Evans et al. [2021]), a model specifically designed for predicting protein complexes. This version represents a more robust and systematic approach, tailored to handle the challenges of multimeric interactions without relying on the improvisations required by earlier methods. Despite these advances, challenges persist, particularly in predicting protein interactions and docking involving multiple structures, as well as in understanding protein dynamics and the relationship between structure and function.

In parallel, other AI-based tools were proposed to estimate/score protein-protein interaction, such DeepPPI Du et al. [2017] which use a fully connected deep neural with relu activation network to process protein sequences and descriptors, GraphPPIS Yuan et al. [2021] which treats proteins as graphs and graph convolutional networks (GCNs) to capture spatial proximity and interface-specific features to predict protein site, PepNN Abdin et al. [2022] that focus on peptide-protein interactions by introducing a reciprocal attention mechanism simultaneously updating embeddings for peptides and proteins, ensuring symmetry and dynamic learning, DeepInteract Morehead et al. [2022] that uses a geometric-aware transformer to predict protein interface contacts and integrated rotational and translational invariance, making it suitable for 3D structure-based predictions, DL-PPI Wu et al. [2023] which incorporated Inception modules and a Feature-Relational Reasoning Network (FRN), PeSTo Krapp et al. [2023] which uses a parameter-free geometric transformer for predicting protein-protein binding interfaces, by directly processing atomic coordinates, TAGPPI (Song et al. [2022]) which combine sequence CNNs with AlphaFold-predicted contact maps, processed through graph neural networks (GNNs).

In this paper however, we investigate specifically on the improvement of AFIG, it is now widely adopted and used (Pacesa et al. [2024], Zambaldi et al. [2024], Wang et al. [2024], Torres et al. [2022], Liu et al. [2024], Zhang et al. [2024], Mergen et al. [2024], Balbi et al. [2024]) and we present several contributions that advance the application of AF2 in this context. Note that very recently, Deepmind released the new Alphafold 3 (AF3) (Abramson et al. [2024]) which enhances AlphaFold 2 on every tasks (monomer and multimer) and that also includes other molecules types (such as RNA, dna, and various non-peptidic ligands). However, fewer works have benchmarked AF3 and AF3 usage is more restricted (see AF3 TERMS OF USE). For that reason, our current work focuses on the use of AF2, releasing to HAPI to provide a more open resource to the community.

First, we introduce HAPI (Hacky API), a Python library designed to streamline the integration of AF2 into computational workflows. HAPI supports both the monomer and multimer versions of AF2 and includes flexible pipelines for handling MSAs and templates, offering the community a practical tool to experiment with AF2 in diverse scenarios. Beyond providing this resource, we extend AF2’s capabilities by adapting its multimer version for peptide-protein interaction detection, updating approaches such as the one explored by AFIG, which relied on the monomer version. By leveraging the multimer extension, we provide a focused evaluation of the performance of this extension, that we call HAPI-ppi, using the curated Propedia database Martins et al. [2023].

Furthermore, we enhance the utility of AF2/HAPI-ppi by developing predictive models trained on datasets derived from Propedia, where AF2’s outputs serve as input features. To this end, we assess classical machine learning models as baselines while introducing PeTriPPI a fine-tuned version of our new PeTriPOV transformer encoder, update from PeTriBERT Dumortier et al. [2022] which is then adapted for this task. Our results demonstrate that this boosting methodology significantly improves prediction performance on our dataset compared to relying solely on AF2’s raw outputs.

Building on previous work and addressing key challenges, our study combines practical tools, methodological updates, and empirical insights to contribute to the ongoing development of peptide-protein interaction prediction.

## 2. Results

### 2.1 AF2 multimer extension for PPI detection and scoring

From AlphaFold 2.3.2, we wrote HAPI python code, and designed HAPI-ppi, which is based on i) a scooped datapipeline where no MSA is computed ii) Set the full pdb atom coordinates as structure template to the AF2 neural network. Unlike AFIG however, we didn’t implemented the structure module initialization and kept the original implementation of AF2 that initialize the residue frames at the global frame. To compare HAPI-ppi to AFIG, HAPI-ppi was run on a supercomputer on a curated version of the propedia dataset, that contains about 10,000 tuples of interacting peptide/protein interactions. To produce negative examples, we followed the assumption of sparsity of the interacting proteins and randomly draw non-existing couples of peptide/protein of the dataset for which we artificially generated pdb files with correct tertiary conformation. Then, we compared HAPI-ppi and AFIG using the PAE interaction scores and the same threshold of 10 from the original paper. The HAPI-ppi estimator, based on the multimer version of AlphaFold, outperformed AFIG across 500 interaction cases (25% positive). As shown in Table 1, HAPI-ppi achieved a precision of 0.88 and an AUC of 0.76, outperforming AFIG’s 0.54 precision and 0.68 AUC. Notably, while recall metrics remained low, HAPI-ppi showed significant precision improvements, marking a significant advancement over AFIG. Interestingly, these results were achieved with HAPI-ppi without employing the geometric frame initialization that assumes the whole quarternary conformation is known and which is used in AFIG’s structure module; providing complex structures as templates to the Evoformer was sufficient for superior interaction classification. This is further illustrated in Figure 1, where HAPI captures more of the interacting cases than the AFIG models. This suggests that the AF2 multimer model may require less contextual information for estimating interactions.

**Table 1:** Results of logistic regression on the PAE interaction scores (25% positive interactions), comparing AlphaFold- HAPI and AFIG with structure module inialization.

| Model | Precision | Recall | Log Loss | AUC ROC |
| --- | --- | --- | --- | --- |
| AlphaFoldHAPI | **0.88** | **0.50** | **0.42** | **0.76** |
| AFIG + structure module initialisation | 0.54 | 0.25 | 0.51 | 0.68 |

**Figure 1:**
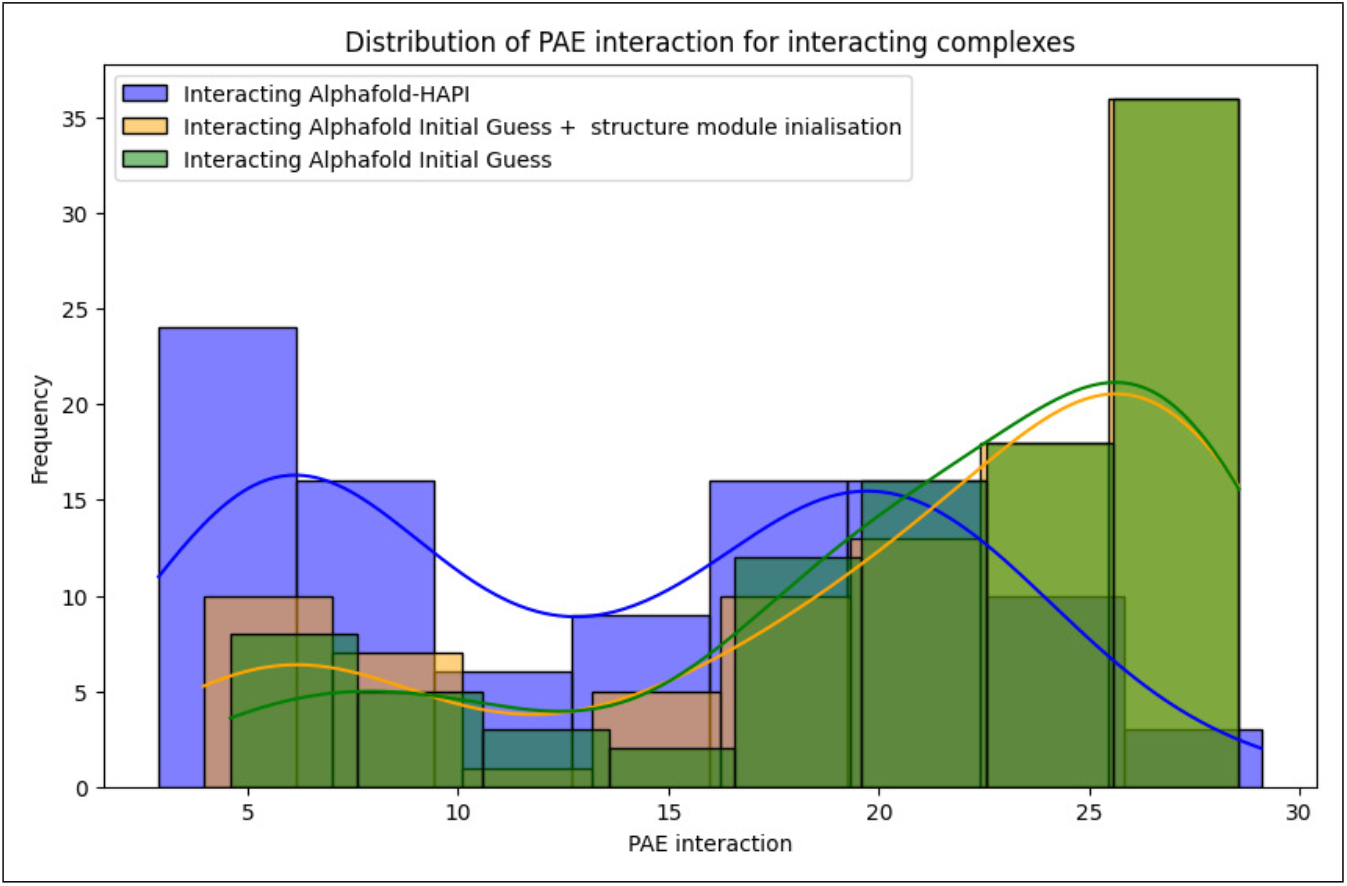
Comparison of real interacting peptide-protein repartition across the pae interaction scores computed with AlphaFold-HAPI, AlphaFold Initial Guess with/without structure module initialization

We also remarquably noted that the empirical PAE interaction distribution of known interacting complexes, as seen in Figure 2, shows a bimodal distribution of both positively interacting and negatively interacting peptide-protein pairs.

**Figure 2:**
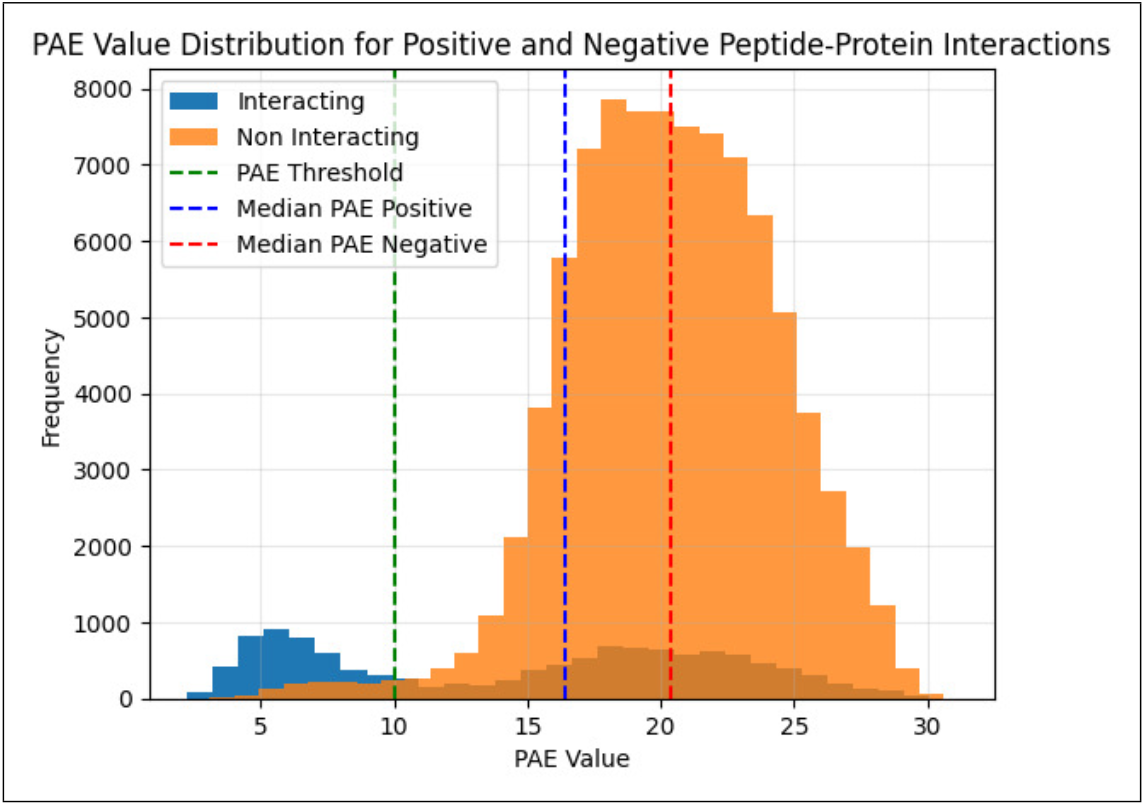
Bimodal distribution of interacting and non-interacting peptide-protein cases along PAE interaction scores obtained from AlphaFold-HAPI

This analysis, conducted on 100,000 cases with 10% positives highlights the asymmetry of the PAE interaction estimator, often used as a threshold in numerous application which shows with very few false positives below the 10 Å threshold set by Bennett et al. [2023]. However, some real positive interactions still remain undetected, pointing to the precision of the method but also emphasizing its recall limitations. Although this approach is useful for distinguishing between interacting and non-interacting peptides, the reliance solely on PAE interaction metrics leaves a considerable proportion of false negatives, raising the question to know wether if AF2 might leave positive examples aside and naturally indicating that further refinement might needed to improve detection accuracy across all cases and further investigation is necessary to interpret this bimodality clearly.

### 2.2 Boosting HAPI with Machine Learning and AI

To enhance the classification power of AlphaFold HAPI, we hypothesized that AF2, while biased toward interacting structures, might still have captured deep interaction-related information in its network which we hoped could be retrieved using advanced pattern recognition techniques.

In particular, HAPI/AF2 outputs include various features known as pLDDT, PAE, IPTM, and PTM, which have varying modalities and spatial associations. These are either spatially confounded with residues (e.g., pLDDT), residue pairs (e.g., PAE), or be global attributes (e.g., PTM, IPTM). The variability in feature dimensions, which correlates with protein size, poses challenges for classical machine learning methods that often require fixed-dimension inputs. Nonetheless, we still explored two distinct approaches: i) moment-based methods, used as baselines that performs regression on various moments of the metrics but which don’t benefit from the spatial information leaving them behind ii) A fine-tuned ad-hoc transformer model that can be fed with the different spatial features.

#### 2.2.1 Moment-Based Methods Classification

Moment-based methods reduce feature complexity by summarizing distributions of AlphaFold-derived metrics, such as pLDDT and PAE, into statistical moments. While these approaches yield high precision, they fail to capture the nuanced spatial interactions critical for improving recall, highlighting the need for advanced models like PeTriPOV and PeTriPPI. For example, pLDDT and PAE were summarized using statistical moments, enabling multivariate analysis on simplified features.

We applied various machine learning models, including Random Forest and XGBoost, to these features in order to predict peptide-protein interaction from HAPI-ppi (AF2) features. Random Forest achieved the highest precision (0.88) and balanced precision-recall with an F1-score of 0.61 (Table 2). However, recall remained low across all models, suggesting that moment-based methods struggle to capture all true positive interactions.

**Table 2:** Performance of machine learning models on interaction prediction.

| Model | Precision | Recall | F1-Score | Total Accuracy | Mean Accuracy (k-fold) |
| --- | --- | --- | --- | --- | --- |
| Logistic Regression | 0.80 | 0.37 | 0.51 | 0.9154 | 0.9148 |
| **Random Forest** | **0.88** | **0.46** | **0.61** | **0.9298** | **0.9282** |
| XGBoost | 0.85 | **0.47** | 0.60 | 0.9154 | 0.9148 |
| SVM (RBF Kernel) | 0.82 | 0.37 | 0.51 |  |  |
| SVM (Linear Kernel) | 0.79 | 0.38 | 0.52 | 0.9160 | 0.9144 |
| LGBM | 0.85 | 0.43 | 0.57 | 0.9237 | 0.9213 |
| CatBoost | 0.86 | 0.45 | 0.59 | 0.9267 | 0.9251 |
| GBM | 0.82 | 0.39 | 0.53 | 0.9188 | 0.9166 |
| AdaBoost | 0.79 | 0.38 | 0.51 | 0.9157 | 0.9143 |

Correlation analysis (Figure 3) revealed strong relationships between key metrics, such as PAE_interaction_mean and plddt_binder, highlighting potential redundancies. PCA was also explored to reduce feature complexity, but retaining the full set of moments proved more effective for downstream models. Adding higher-order moments, such as skewness and kurtosis, marginally improved some models but did not consistently enhance performance.

**Figure 3:**
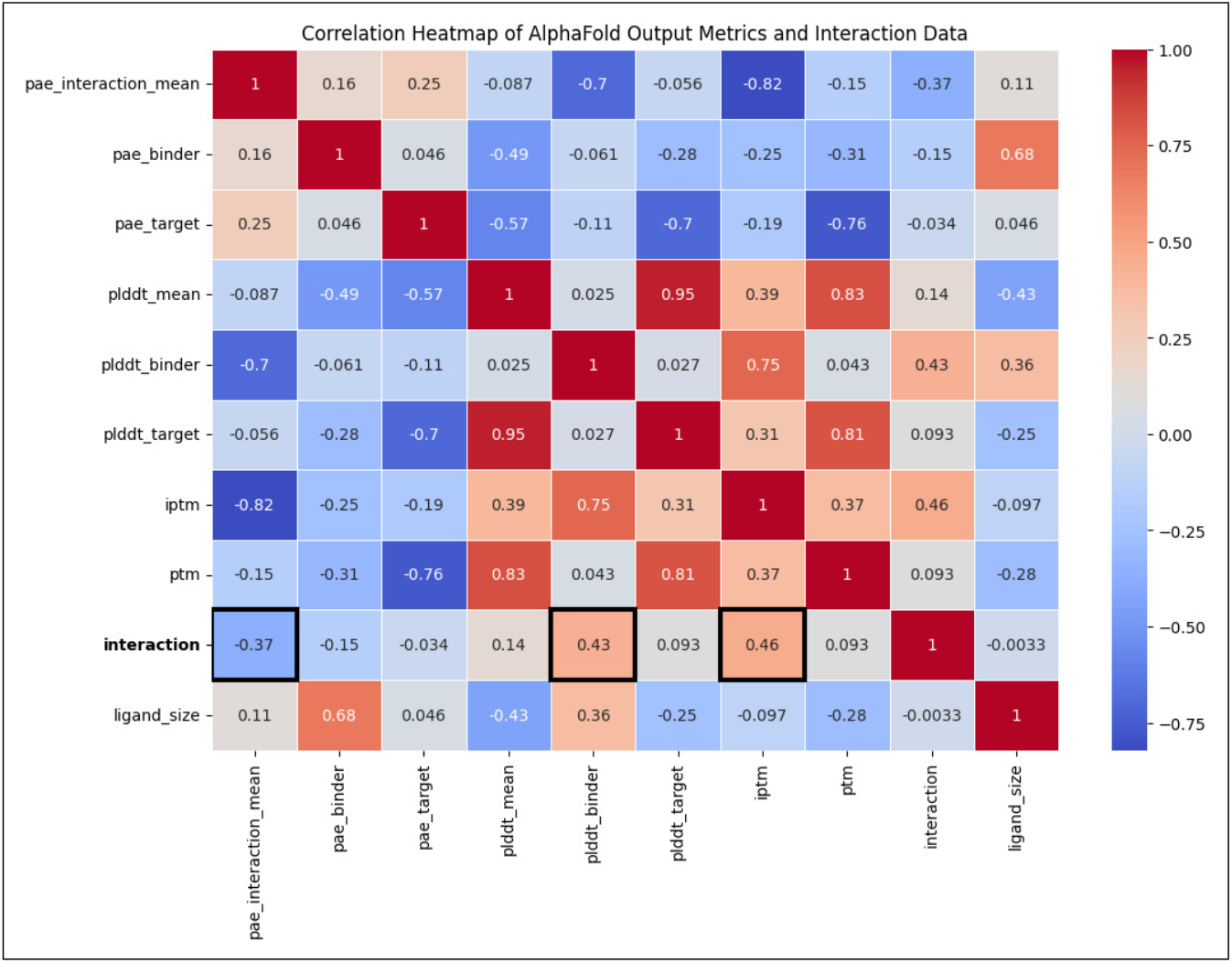
Heatmap showing correlations between AF-HAPI-ppi output metrics and interaction data. Strong correlations are observed between interaction status and metrics like PAE_interaction_mean and plddt_binder.

**Figure 4:**
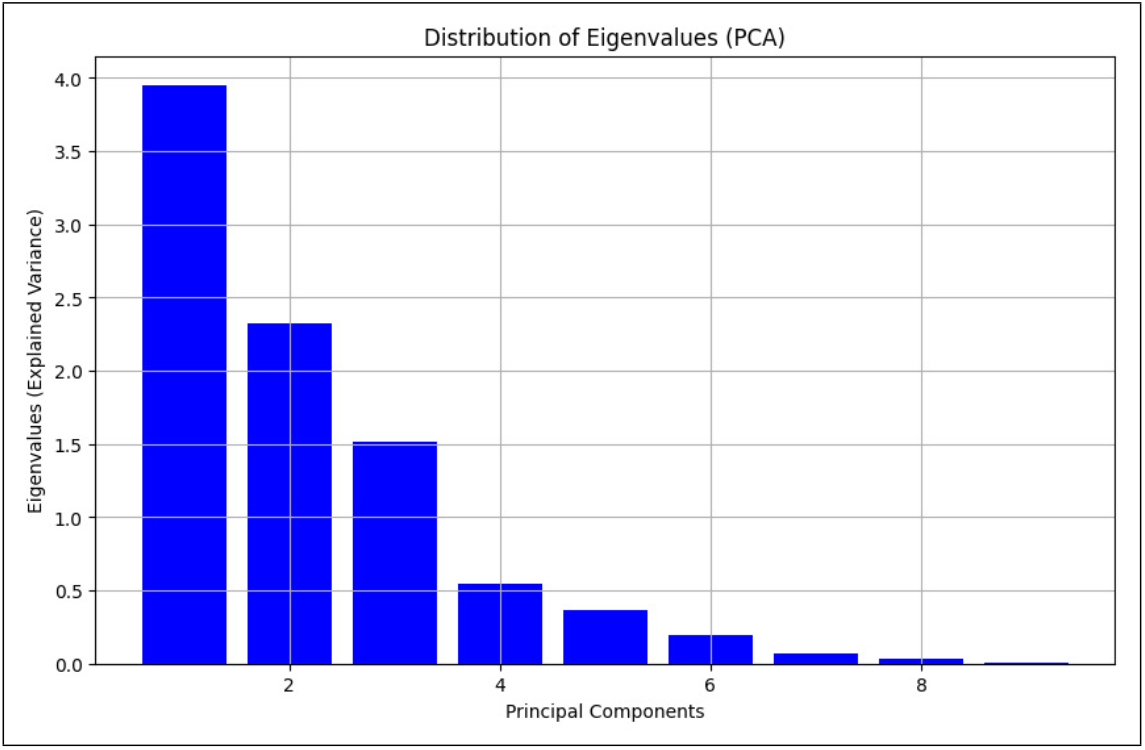
Variance explained by the first four PCA components of AF-HAPI-ppi features.

While precision was high, the low recall suggested that critical interaction patterns were not fully captured by moment-based methods. The bimodal distribution of PAE, coupled with limitations in other metrics, indicated that spatial data might hold additional information critical for improving recall.

#### 2.2.2 PeTriPOV Transformer Encoder

To develop an effective PPI detector, we fine-tuned a novel transformer-based encoder designed to mitigate overfitting on the training dataset. This encoder builds upon our prior work (Dumortier et al. [2022]), which introduced a transformer trained through Masked Language Modeling (MLM) on sequences composed of the 20 standard amino acids in proteins. While this approach captures sequence information, our earlier model also incorporated structural features. However, PeTriPOV extends this capability by introducing a refined mechanism for encoding structural relationships that enhances both training performance and generalization.

The PeTriPOV model differs fundamentally from its predecessor, PeTriBERT, in how structural information is encoded. PeTriBERT employs an absolute encoding of position and rotation within the sequence vectors. In contrast, PeTriPOV encodes structural features by leveraging relative transformations—specifically, differences in rotation and translation between consecutive residues. Relative transformations between residues are featurized by a lightweight MLP, producing NxN vectors that encode pairwise relationships. These vectors are added as a bias term in the transformer’s attention mechanism, dynamically integrating spatial context throughout training.

To ensure a consistent basis for comparison, PeTriPOV was trained using the same dataset and hyperparameters as PeTriBERT. The results reveal clear improvements in both training and generalization performance. Specifically, PeTriPOV demonstrates superior fitting on the training dataset, as illustrated in Figure 5b. Notably, despite its enhanced capacity to model the training data, PeTriPOV avoids overfitting, achieving significantly better generalization on both validation and test datasets. Validation loss and validation accuracy across steps are shown in Figure 5c and 5a, highlighting the consistent improvements across epochs. The test dataset results, summarized in Table 3, further confirm the superior performance of PeTriPOV in comparison to PeTriBERT.

**Figure 5:**
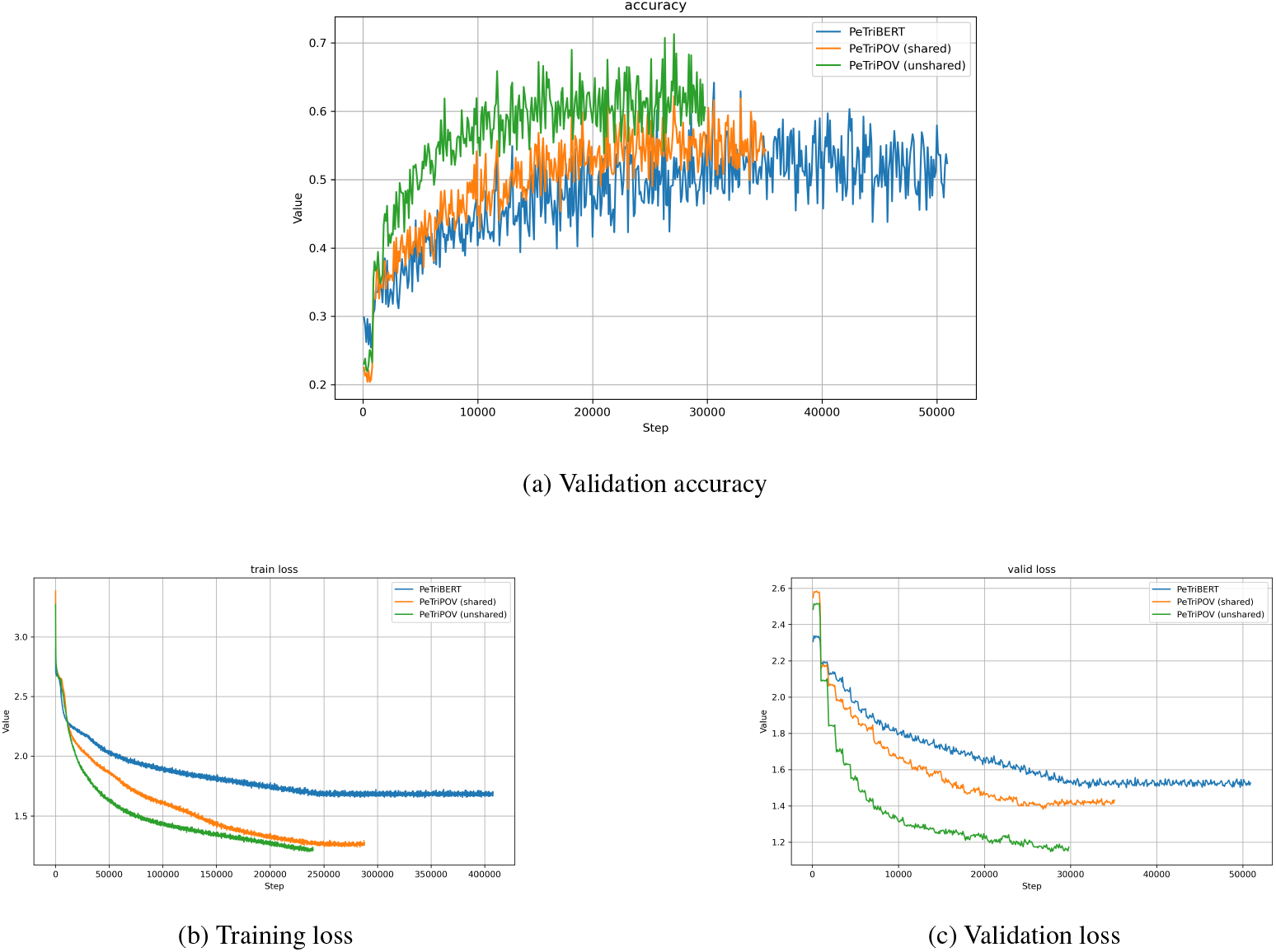
Results of our experiments. The figure shows the performance of the three models—PeTriBERT, PeTriPOV (shared weights), and PeTriMPOV (unshared weights)—across different metrics: validation accuracy, training loss, and validation loss. PeTriPOV (shared and unshared) consistently outperform PeTriBERT, showcasing better generalization and training efficiency. The validation and training loss curves also demonstrate the robust convergence of our models. Furthermore, the version with unshared weights at each layer (PeTriMPOV) achieves better overall performance without overfitting, highlighting the importance of independent layers in improving model expressiveness and generalization.

**Table 3:** Performance metrics for different models.

| Model | Cross Entropy | Accuracy | Precision | Recall | F1 |
| --- | --- | --- | --- | --- | --- |
| PeTriBERT | 1.54 | 0.52 | 0.53 | 0.52 | 0.51 |
| PeTriPOV (shared weights) | 1.42 | 0.55 | 0.55 | 0.55 | 0.54 |
| PeTriPOV (unshared weights) | **1.25** | **0.60** | **0.60** | **0.60** | **0.59** |

The improved generalization of PeTriPOV can be attributed to its ability to encode structural information more effectively. By incorporating relative differences in rotation and translation directly into the attention mechanism, the model gains a nuanced understanding of protein structural dynamics. This design enables PeTriPOV to capture complex structural relationships critical for PPI prediction, while maintaining robustness across diverse datasets. These results underscore the utility of relative structural encoding as a pivotal improvement over absolute positional representations in transformer-based architectures for protein sequence and structure analysis.

#### 2.2.3 PeTriPPI Transformer

Moment-based methods often lose spatial information about feature positions within the sequence, potentially discarding patterns encoded by AlphaFold that are critical for predicting interactions. To overcome this limitation, we developed PeTriPPI, a transformer-based model specifically fine-tuned for PPI detection. PeTriPPI preserves both sequence and structural information, enabling a more comprehensive understanding of peptide-protein interactions compared to moment-based approaches.

Building on a pretrained PeTriPOV encoder, PeTriPPI incorporates additional modifications tailored to the PPI task. The model employs a task-specific head for PPI detection, replacing the standard transformer output layer, and leverages AlphaFold’s high-accuracy protein interaction (HAPI) features. Features such as pLDDT and PAE are featurized via adapter layers and seamlessly integrated into the encoder, enriching its representation of spatial relationships critical for interaction detection. By leveraging AlphaFold features such as pLDDT and PAE within adapter layers, PeTriPPI enriches its spatial representation, leading to a notable improvement in recall metrics, as shown in Table 4. This demonstrates the advantage of combining pretrained structural embeddings with task-specific fine-tuning for PPI detection.

**Table 4:** Results of the fine-tuned PeTriPPI model on the PPI task using the Propedia dataset.

| Model | Precision | Recall | F1 Score |
| --- | --- | --- | --- |
| PeTriPPI | 0.83 | 0.74 | 0.71 |

To minimize overfitting during fine-tuning, we adopted a multi-faceted approach. A low learning rate (1e-5), gradient boosting, and a weighted cross-entropy loss function were employed to address class imbalance. Additionally, freezing the pre-trained transformer parameters during the initial training phase further mitigated the risk of overfitting, ensuring stable and efficient learning.

The results, presented in Table 4, underscore the advantages of incorporating both sequence and structural features. PeTriPPI achieved a recall of 0.74, surpassing the baseline machine learning models in this metric. While precision showed marginal improvement compared to moment-based methods, the significant boost in recall highlights the model’s ability to capture subtle patterns indicative of interactions, thereby improving overall robustness in PPI detection.

By effectively integrating sequence and structural features derived from AlphaFold, PeTriPPI addresses the key limitations of moment-based methods. The results demonstrate that preserving structural context is crucial for accurately capturing complex interaction patterns, paving the way for more reliable and interpretable PPI predictions.

## 3 Discussion

This study highlights the significant strides made by the multimer version of AlphaFold in advancing our understanding of peptide-protein interactions. To summarize, the core ideas of our approach are: i) the acceleration of AlphaFold’s data pipeline without sacrificing too much performance, ii) the use of AlphaFold multimer and iii) the introduction of PeTriPPI, a fine-tuned transformer-based model that builds upon PeTriPOV. PeTriPPI leverages the pretrained embeddings of PeTriPOV but goes further by introducing additional types of embeddings, modifying the decoder layer, and retraining the primary encoder layers to better capture peptide-protein interaction-specific features. This iterative improvement allows PeTriPPI to deliver enhanced predictions tailored to the complexities of peptide-protein interactions.

Notably, even when dropping the learning-based components and solely using PAE interaction features, HAPI retains strong precision (0.83) compared to previous AlphaFold Initial Guess (AFIG), as it benefits from the multimer submodel of AlphaFold. This highlights the robust foundation provided by AlphaFold-derived features in peptide-protein interaction prediction workflows.

Additionally, PeTriPOV itself represents a significant improvement over our earlier model, PeTriBERT, by incorporating independent layer structures that better capture spatial and sequential features of peptide-protein interactions. PeTriPOV serves as a robust pretrained encoder, laying the foundation for more specialized models like PeTriPPI. Integrating PeTriPOV as a pretrained encoder into our workflow has yielded further performance improvements. By fine-tuning PeTriPPI on peptide-protein specific tasks, we effectively harness AlphaFold’s features in novel ways, boosting the model’s effectiveness in identifying complex interactions. By comparing PeTriPPI with traditional machine learning baselines, we show that our approach offers substantial advantages, making the baseline methods less relevant for advanced tasks. The ability of PeTriPPI to retain both sequence and structural information represents a significant leap forward in PPI prediction. This advancement not only improves computational efficiency but also lays the groundwork for more accurate in silico screening of peptide-based therapeutics. Future extensions could explore fine-tuning PeTriPOV on larger, multi-modal datasets to further enhance its applicability to diverse biological systems.

Our work contributes to improving the filtering steps of peptide-protein designs performed in in silico experiments before transitioning to more expensive in vivo testing. Given the significantly lower costs of in silico experiments, the trade-off between high precision and lower recall is acceptable. By prioritizing precision, we ensure that the selected designs are those with the highest confidence, reducing the likelihood of false positives and streamlining subsequent experimental workflows.

Finally, we hope this study, alongside the HAPI code we provide, contributes meaningfully to the community. HAPI is both a toolbox that enables easier integration of AlphaFold into Python workflows—by decoupling its data pipeline from its neural network—and a filtering method for peptide-protein interaction prediction, HAPI-ppi. By sharing these tools, we aim to facilitate future research in peptide design and protein interaction studies, providing an accessible and customizable framework for further exploration. We believe that the introduction of PeTriPOV and PeTriPPI paves the way for the development of new drugs or agrochemicals, demonstrating the promise of transformer-based models in this critical domain.

## 4 Methods

### 4.1 HAPI

AlphaFold2 (AF2) revolutionized protein structure prediction by employing a deep learning architecture that integrates evolutionary information from multiple sequence alignments (MSAs) and structural templates. While the neural networks operate efficiently during inference—typically benefiting from GPU acceleration—the data pipeline is comprised of a series of clustering tools such as JackHMMER, HHblits, or HHsearch (Johnson et al. [2010], Remmert et al. [2012], Steinegger et al. [2019], Eddy [2011]), which can significantly delay processing times. Numerous studies have sought to expedite this process, either by accelerating the construction of MSAs Mirdita et al. [2022] or by substituting MSA computation with alternative deep learning layers Lin et al. [2022], Chowdhury et al. [2022]. However, these investigations have consistently confirmed that performance drops when MSAs are not incorporated.

To address these challenges and improve the usability of AF2 within Python, we developed HAPI (Hacky API). HAPI encapsulates the original AF2 functionalities in an object-oriented framework, enabling researchers to seamlessly integrate AF2 into Python-based workflows. The primary interface in HAPI is the “AlphaFolder” object, which accepts sequence strings or file inputs (e.g., FASTA or PDB files) and provides streamlined methods for feature computation and structure prediction (see Figure 6).

**Figure 6:**
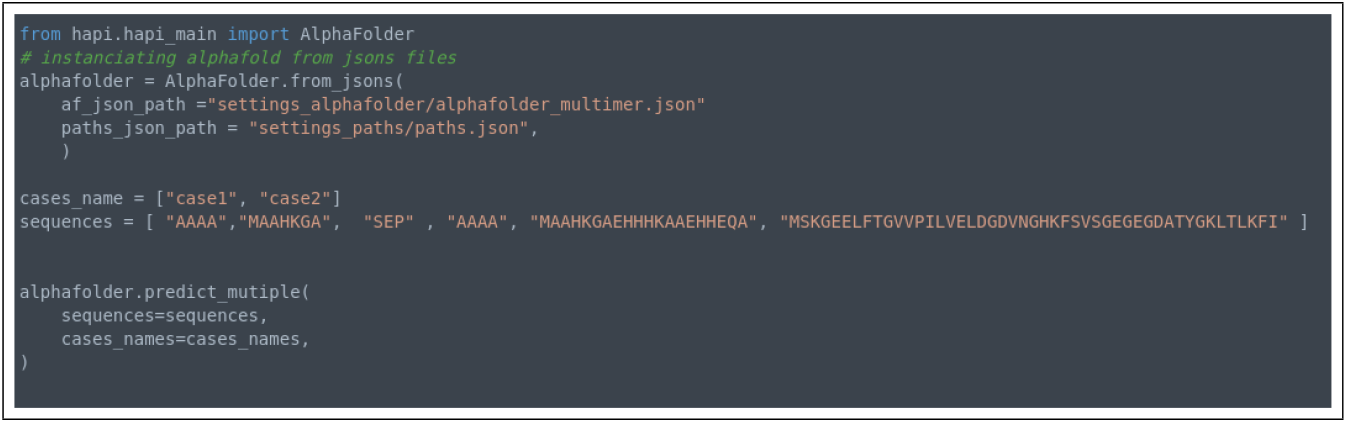
Snippet of use of HAPI.

HAPI also decouples the data pipeline from the neural network component, providing flexibility to experiment with alternative configurations. Users can switch between multiple data pipeline modes, including:

- Standard mode: computing MSAs and templates for monomers.
- Single-sequence mode: bypassing MSA and template computation.
- Monomer mode with a PDB file as a template (no MSA computation).
- Multimer mode with a PDB file as a template (no MSA computation).
- Dimer mode with a PDB template and MSA computation for one chain (ligand-receptor mode).

This modular approach not only simplifies the customization of AF2 workflows but also enables advanced use cases, such as ligand-receptor modeling or peptide-protein interaction detection. HAPI will be made freely available to the research community at https://github.com/Baldwin-disso/Alphafold2_HAPI.

### 4.2 HAPI-ppi

While the original AF2 implementation excels in monomeric structure prediction, it has also been shown to be effective for protein-protein interaction (PPI) analysis. Studies such as Bryant et al. [2022a] and Bennett et al. [2023] demonstrated that AF2 can predict interactions by using confidence metrics derived from its output, such as the predicted local distance difference test (pLDDT) and predicted aligned error (PAE) matrices. Bennett et al. Bennett et al. [2023] identified two key failure modes for binders (peptide ligands): i) the binder fails to adopt the intended structure, and ii) the binder fails to bind to the target receptor. They proposed a dual scoring criterion to select effective designs, based on pLDDT and PAE values, which was shown to improve experimental outcomes.

Building on these insights, HAPI-ppi adapts AF2 for peptide-protein interaction detection by leveraging the multimer model and simplifying the data pipeline in a similar way than AFIG. HAPI-ppi streamlines the AF2 pipeline by bypassing database searches and using ground-truth structures as templates, significantly reducing runtime. In this configuration, the input to the Evoformer is a single target sequence, while the structural templates are replaced with the complex structure for which the interaction is being tested. This simplified scheme is illustrated in Figure 7.

**Figure 7:**
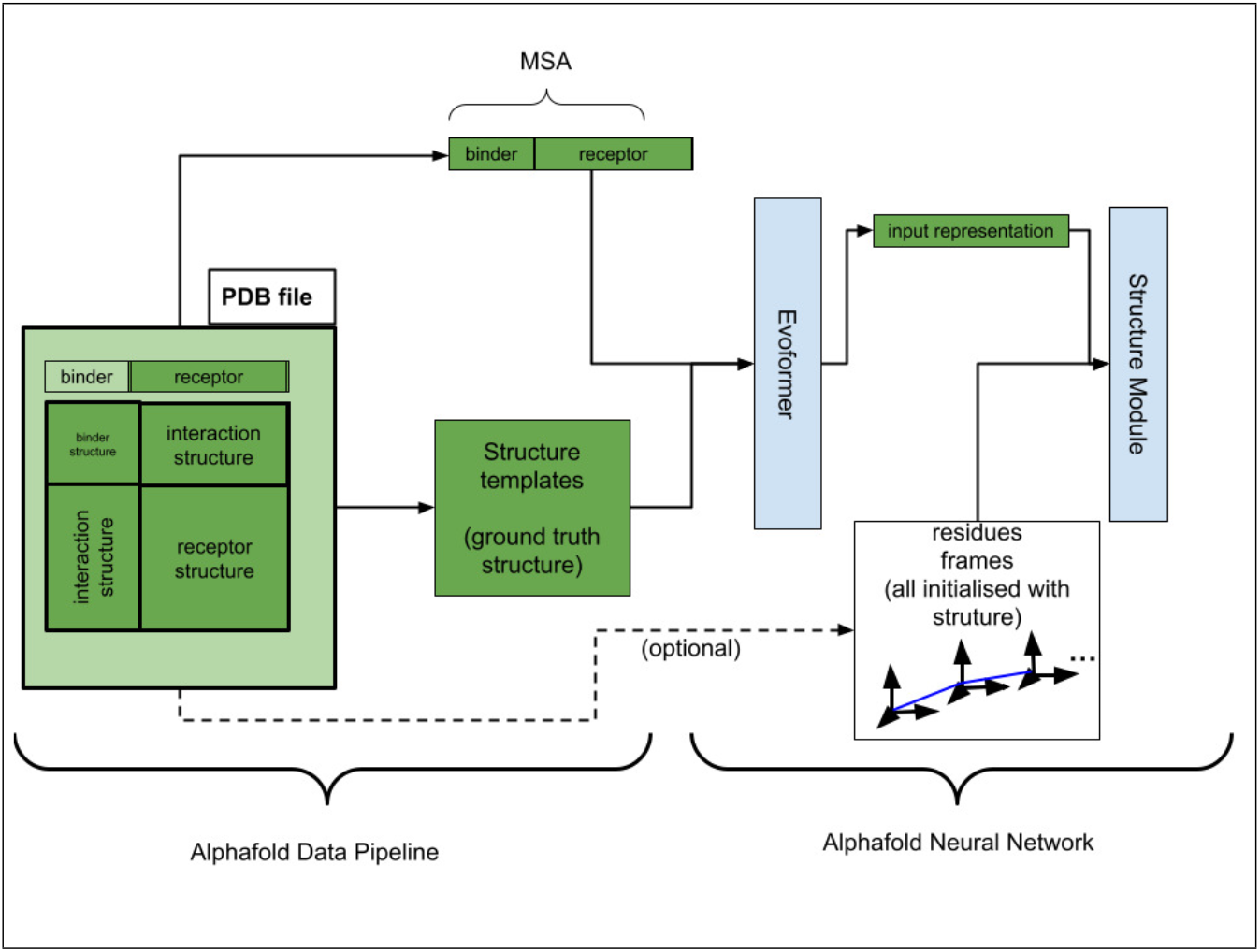
Global scheme of HAPI-ppi (inspired by AlphaFold Initial Guess Bennett et al. [2023]).

By leveraging these adjustments and building upon prior methodologies, HAPI-ppi provides a flexible framework for detecting PPIs while incorporating additional strategies to improve performance, which will be detailed in subsequent sections of this article.

### 4.3 Dataset and Data Augmentation

We evaluated our methodology using the Propedia Dataset v2.3 Martins et al. [2023], which comprises approximately 20,000 experimentally validated models of peptide-protein interactions in PDB format. To augment this dataset, we randomly sampled peptide-protein pairs, assuming a sparsity of interactions (interactions absent from the dataset are unlikely to occur), as posited by several well cited works in biology (Yu et al. [2008], Rolland et al. [2014], Venkatesan et al. [2009]). This process yielded approximately 100,000 samples, with 10% categorized as positive cases. Utilizing the computing cluster Jean Zay (http://www.idris.fr/eng/jean-zay/), we processed the entire augmented dataset with HAPI-ppi, documenting all output features and totalling 1.7 TB of feature data. We employed a simplified data pipeline that did not compute MSAs and utilized provided .PDB structure files as templates for both ligands and receptors.

### 4.4 HAPI/AF2 output features

To further analyze the data, we extended our approach by applying classical machine learning techniques to the features derived from AF-HAPI outputs:

- pLDDT (predicted Local Distance Difference Test (Mariani et al. [2013])) : The pLDDT (predicted Local Distance Difference Test) is a confidence metric used by AF2 to evaluate the local accuracy of its predicted protein structures. It provides a measure of how reliable the prediction is for each region or residue within a protein. This score is crucial for understanding which parts of a predicted structure are likely to be correct and which should be treated with caution. pLDDT is residue-specific : it provides a per-residue score, allowing scientists to visualize which parts of the protein are well-predicted (high pLDDT) and which parts might have inaccuracies or flexibility (low pLDDT). it ranges from 0 to 100: a score above 90 indicates high confidence in the local structure, meaning AF2 predicts that region with high accuracy, a score between 70 and 90 suggests moderate confidence, and scores below 50 indicate low confidence, meaning that part of the protein might be unreliable or incorrectly modeled.
- PAE (Predicted Aligned Error (Jumper et al. [2021])) : The PAE (Predicted Aligned Error) is another confidence metric used by AF2 to evaluate the global accuracy of a predicted protein structure, particularly focusing on the relative positions of residues or domains within the protein. While pLDDT measures local confidence for individual residues, PAE provides information about how accurately AF2 predicts the spatial relationships between different parts of the protein. PAE estimates the error or uncertainty in the relative position of two residues or regions in a protein. It assesses how accurately AF2 predicts the alignment between different parts of the structure. It is expressed in angstroms (Å) and indicates the predicted alignment error between two residues or regions. A low PAE score (close to 0 Å) means that AF2 predicts the relative positioning between those residues with high accuracy while a high PAE score means there is significant uncertainty in how those parts of the protein are positioned relative to each other.
- pTM (predicted TM-score) : The pTM (predicted TM-score) is a confidence metric used by AF2 to assess the global accuracy of its predicted protein structures. It is an adaptation of the traditional TM-score, which measures how similar a predicted protein structure is to its true, experimentally determined structure. The pTM score specifically estimates the reliability of AF2’s predictions regarding the overall three-dimensional fold of a protein.
- ipTM (inter-chain predicted TM-score) : The ipTM (interface predicted TM-score) is a metric developed by AF2 to specifically assess the quality of protein-protein interfaces in multi-protein complexes. It is an extension of the pTM score, but it focuses on how well AF2 predicts the interaction surfaces between different proteins or subunits in a complex.

These features are output by AF-HAPI and stored in .pkl files that will then be further processed.

Indeed, PAE and pLDDT scores are the most important metrics used as classification features. In particular, in the case AlphaFoldInitial Guess, part of the data provided by AF2 were found to be correlated to interaction or stability :

- pLDDT vector can be partionned into two parts : pLDDT binder and pLDDT target that respectively describe mean local folding quality. In the work of Bennett et al. [2023].
- PAE interaction can be partitionned in 3 parts : PAE binder, PAE target and PAE interaction (as illustrated on figure 8). In the work of Bennett et al. [2023], PAE interaction is the main estimator for peptide-protein interaction.

**Figure 8:**
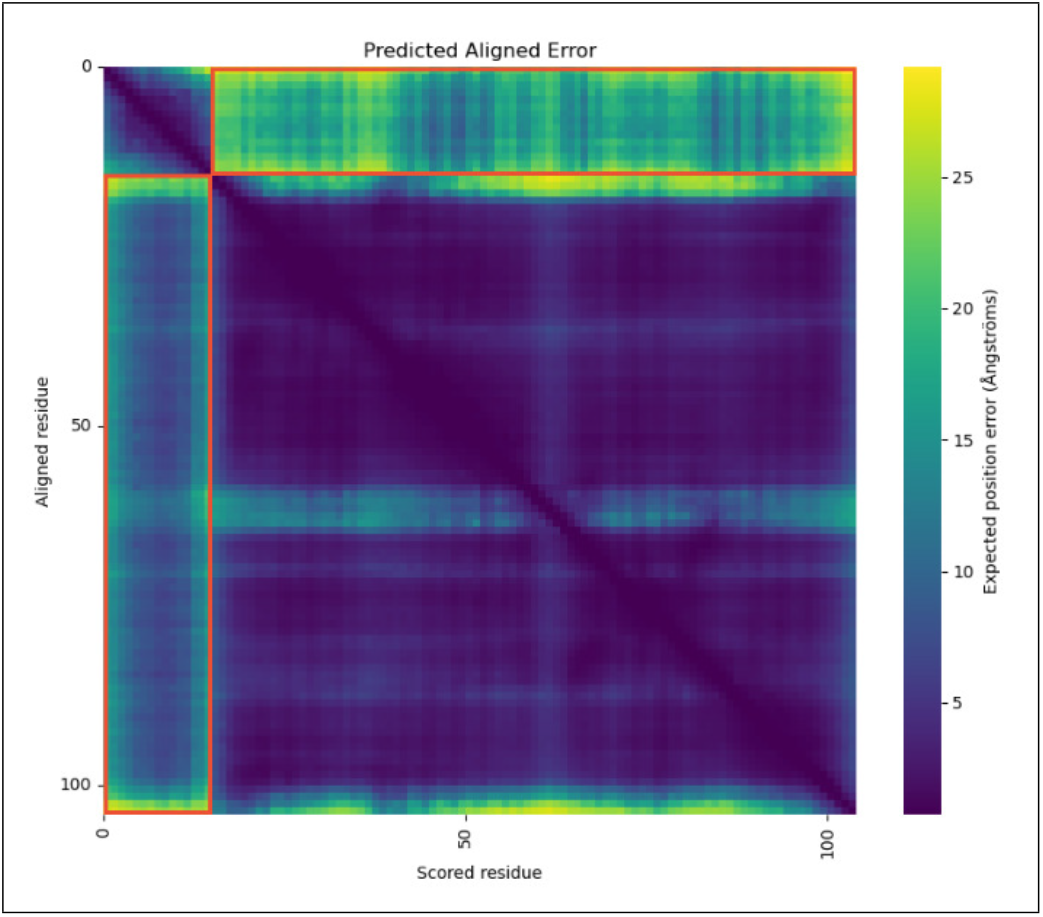
Example of PAE score matrix output by AF2 on a ligand-receptor example. The Upper left diagonal blocs is the part of the PAE matrix where values represent the error estimated for the ligand relatively to itself. Similarly, the inferior right part of the matrix is the part of the PAE matrix associated to the receptor with itself. Finally the antidiagonal blocs (framed with red rectangles) represent the error of the ligand to the receptor. The average of the contents of these red frames is called PAE interaction and is used as the primary features that indicates if an interaction exists or not, by AFIG and our method.

### 4.5 Baseline Machine Learning Methods

#### Linear Models

Linear models, such as Logistic Regression Cox [1958], are used for their simplicity and interpretability. Support Vector Machines (Linear) Boser et al. [1992] extend this by finding a hyperplane that maximizes the margin between classes, improving robustness for linearly separable datasets.

#### Kernel-Based Methods

Kernel methods, such as Support Vector Machines with an RBF kernel Cortes [1995], transform input features into higher-dimensional spaces to handle non-linear separability, enabling more flexible decision boundaries.

#### Ensemble and Boosting Methods

Ensemble methods, including Random Forest Breiman [2001] and Gradient Boosting Machines (GBM) Friedman [2001], combine multiple weak learners to enhance predictive performance. Optimized variants like XGBoost Chen and Guestrin [2016], LightGBM Ke et al. [2017], and CatBoost Prokhorenkova et al. [2018] excel at handling complex data. These methods reduce bias and variance, making them effective for tasks with intricate feature interactions.

#### 4.5.1 PeTriPOV Protein Encoder

Here we present PeTriPOV, a translation and rotation-invariant extension of PeTriBERT Dumortier et al. [2022], derived from a BERT encoder model Devlin et al. [2018] and augmented with protein backbone structure features. PeTriPOV incorporates structural information inspired by AF2 Jumper et al. [2021] and the Structure Transformer Ingraham et al. [2019].

PeTriPOV is a five-layer model with 40 million parameters, differing from PeTriBERT in how it encodes structural information. Similar to AF2, the model defines an orthonormal affine basis for each residue *i* using the Gram-Schmidt process applied to the triangle formed by the *C*_*α*_, *C*, and *N* atoms. A rigid transformation *T*_*i*_ is then used to align the global orthonormal basis with the local residue basis. In PeTriPOV, we encode the relative transformation between two residues *i* and *j*, denoted *T*_*i,j*_, which is computed as:

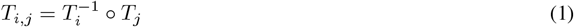

This expression is mathematical rephrasing the the results from the work of (Ingraham et al. [2019]) which has shown this transformation has the desired property to be invariant to global rigid transformation, improves generalization and removes the need to use data augmentation during training. This can trivially trivially proved by the following equation, assuming any global rigid transformation was applied to every residue frames:

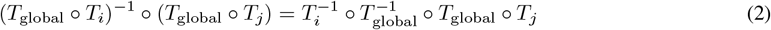

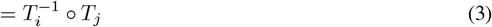

Practically, this transformation is expressed using the centroid **c**_*i,j*_ and rotation matrix *R*_*i,j*_:

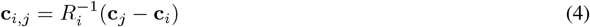

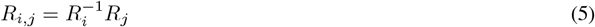

The coefficients of the rotation matrix are compressed into Euler angles, resulting in six parameters (centroid and angles). These parameters are processed by a small fully connected layer to compute a bias matrix **B**. In the transformer, this matrix bias is then added to the attention scores in the attention mechanism:

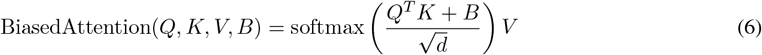

This featurization and attention bias is illustrated on figure 9.

**Figure 9:**
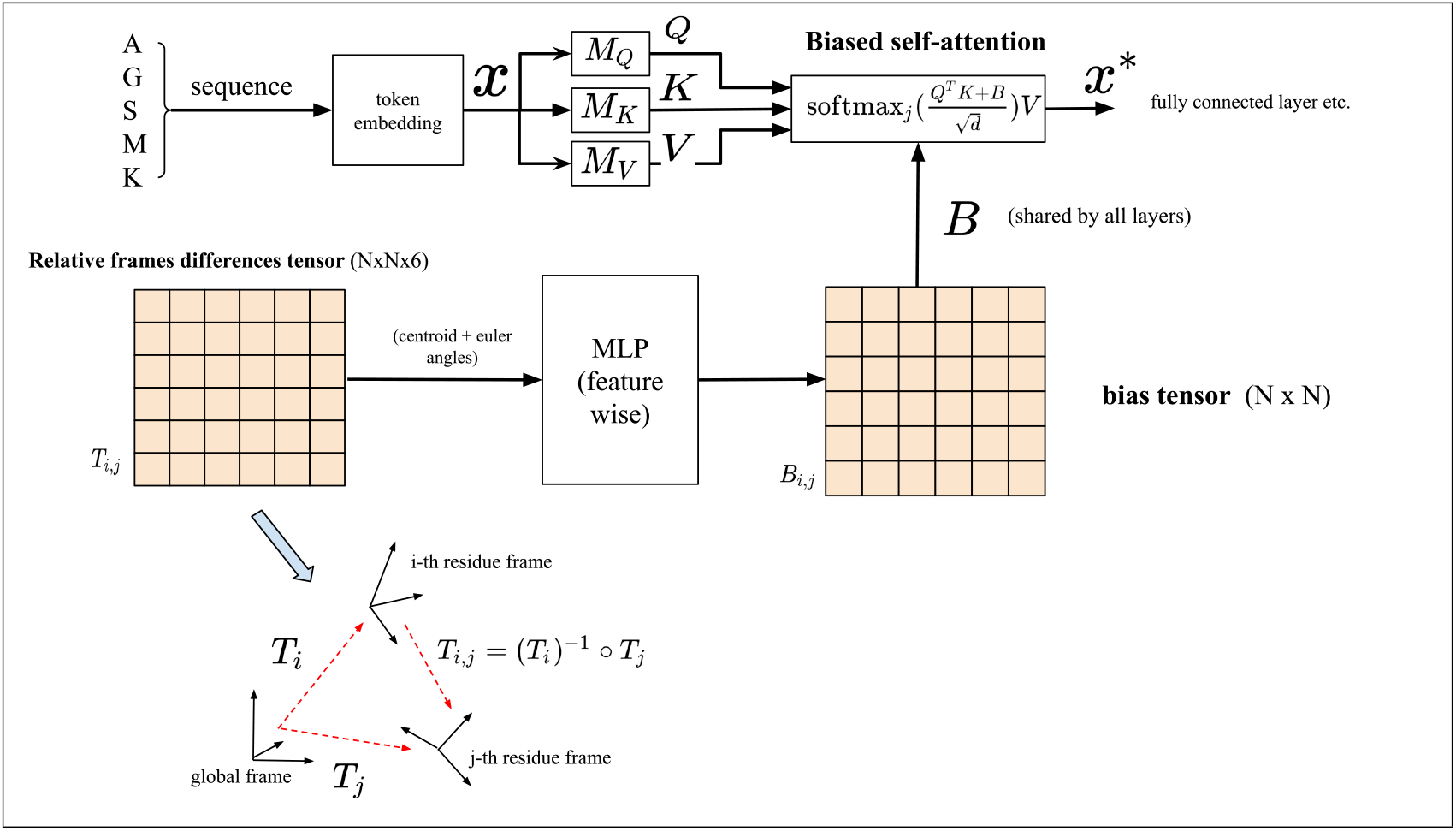
Scheme of the featurization of residue frames and biased attention mechanism in PeTriPOV. A matrix of frames differences is computed by applying to each transformation *T*_*j*_ associated to residue frame *j* the inverse transformation (*T*_*i*_)^*−*1^ to every other residue *i*. Centroids and euler angles parameters of the resulting *T*_*i,j*_ transformations are then passed to an MLP Layer to form a bias Matrix *B*. The bias matrix is different for each attention head but shared amongst layers of the transformer.

This biased attention mechanism is similar to the row-wise gated self-attention with pair bias in AF2. PeTriPOV is trained on approximately 1 million proteins from the AlphaFold Database Varadi et al. [2021] using Masked Language Modeling. The optimization employs AdamW with a learning rate scheduler and warmup.

#### 4.5.2 PeTriPPI: Fine-Tuning PeTriPOV for PPI Detection

For peptide-protein interaction (PPI) detection, PeTriPOV is fine-tuned into PeTriPPI. The fine-tuning process follows approaches similar to those in the original BERT paper Devlin et al. [2018], replacing the model’s classification head with a task-specific, randomly initialized head. This method relates to works such as Rives et al. [2021] and Nambiar et al. [2020] in protein-related tasks.

To incorporate AlphaFold outputs into PeTriPPI, additional feature extraction layers are added, as illustrated in Figure 10. PeTriPPI accommodates sequence-wise (L *×* d) input features by summing them with raw transformer inputs and pairwise (L *×* L *×* d) features as attention biases, added to the *B* matrix. Table 5 details the featurization methods and input types used during fine-tuning.

**Figure 10:**
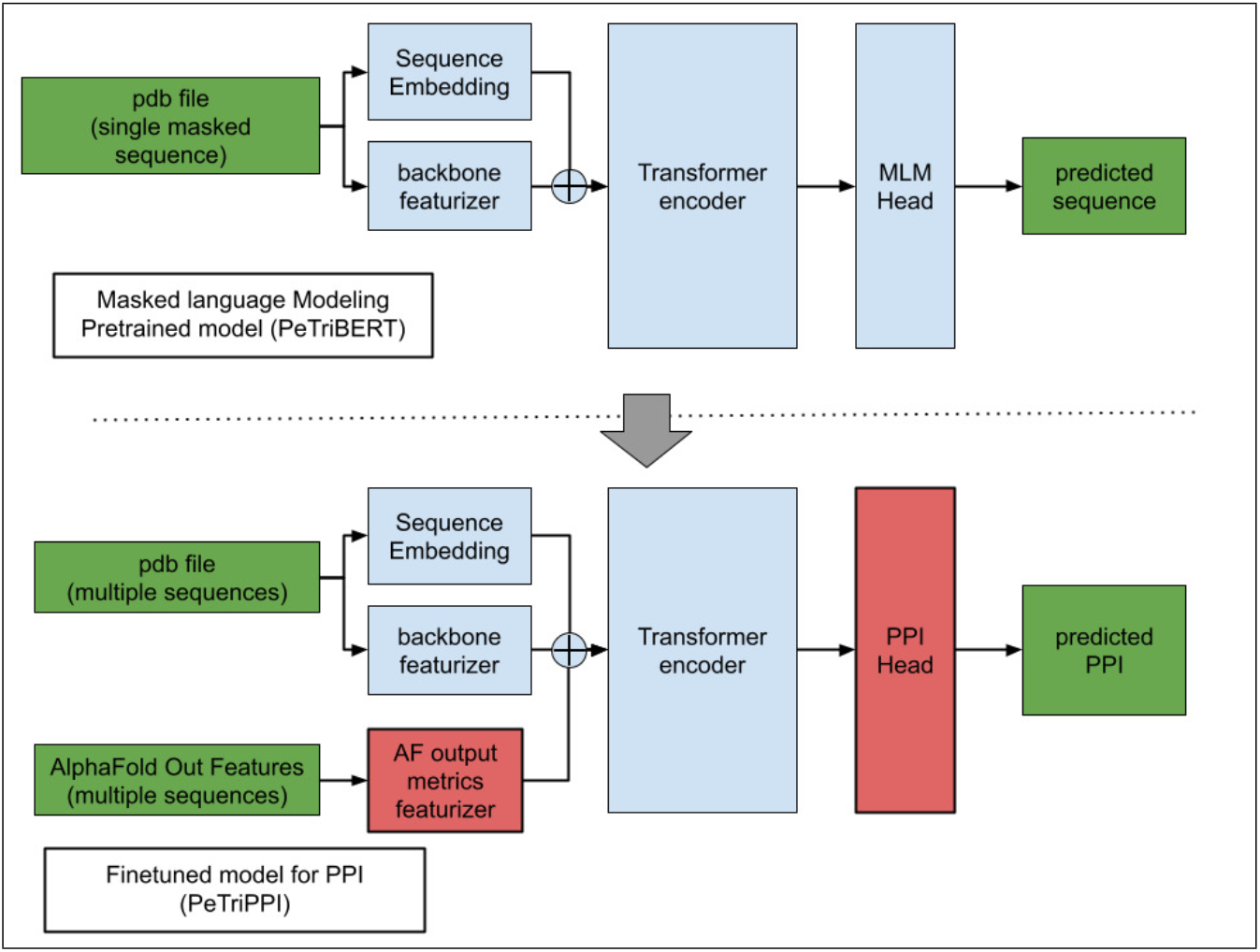
PeTriPOV architecture (top panel) and fine-tuned model for PPI detection (bottom panel).

**Table 5:**
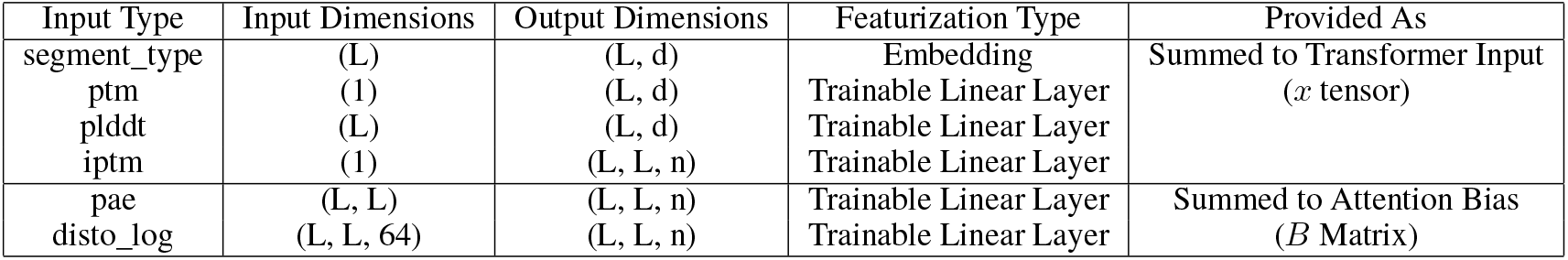
Featurization methods for PeTriPPI inputs.

## 5 Key Points

- We extended the AF2 multimer version to serve as a rapid PPI detection method, demonstrating its superior performance compared to previous methods based on the monomer version of AF2.
- We developed AlphaFold HAPI, a simple Python script that enables convenient integration of AlphaFold within Python code, allowing seamless switching between different data pipelines.
- We further enhanced our approach by applying machine learning to AlphaFold’s outputs to improve detection accuracy.
- Specifically, we fine-tuned an inverse folding deep learning model (PeTriPPI), which proved to be the only method capable of improving the recall of the positive detection class.

## 6 Aknowledgments

Funding: Our work is supported by the CNRS (IRP-CoopNet2), the CNRS innovation program named PEPIA, and by Montpellier University (Isite MUSE project AI3P) to G.K. This work was also publicly funded through ANR (the French National Research Agency) under the program named DeepPep ANR-23-CE20-0020-01.

This work was granted access to the HPC resources of IDRIS under the allocation 2023-AD010714598 made by GENCI.

## 7 Data and code availability

- This work is based on the Propedia 2.3 database (Martins et al. [2023]) available at http://bioinfo.dcc.ufmg.br/propedia2/
- AlphaFoldHAPI and HAPI-ppi are available at https://github.com/Baldwin-disso/Alphafold2_HAPI.
- PeTriPPI graph model are available at https://github.com/Baldwin-disso/PeTriBox.

